# Bat-fruit networks structure resist habitat modification but species roles change in the most transformed habitats

**DOI:** 10.1101/554246

**Authors:** John Harold Castaño, Jaime Andrés Carranza-Quiceno, Jairo Pérez-Torres

**Affiliations:** Grupo de investigación en Biología de la Conservación y Biotecnología, Universidad de Santa Rosa de Cabal, Santa Rosa de Cabal, Colombia.; Laboratorio de Ecología Funcional, Unidad de Ecología y Sistemática (UNESIS), Departamento de Biología, Facultad de Ciencias, Pontificia Universidad Javeriana, Bogotá, Colombia.

**Keywords:** Coffee Cultural Landscape of Colombia, Complex networks, Chiropterocorous plants, Ecological networks, Phyllostomids.

## Abstract

Species do not function as isolated entities, rather they are organized in complex networks of interactions. These networks develop the ecological processes that provide ecosystem services for human societies. Understanding the causes and consequences of changes in ecological networks due to landscape modification would allow us to understand the consequences of ecological processes. However, there is still theoretical controversy and few empirical data on the effects of network characteristics on the loss of natural environments. We investigate how bat–fruit networks respond to three landscapes representing the gradient of modification from pre-montane forest to a heterogeneous agricultural landscape in the Colombian Andes (continuous forests, forest fragments, and crops). We found that forest contained smaller bat–fruit networks than forest fragments and crops. Modified landscapes had similar ecological network structures to forest (nestedness and modularity), but crops contained less specialized networks compared to forests and fragments and the species role in these habitats change. The networks in the rural coffee landscape maintain their structure in the different transformation scenarios, indicating that seed dispersal services are maintained even in the most transformed scenarios. This could be related to the high heterogeneity present in this rural landscape. Although the number of species does not decrease due to transformations, species change their roles in the most transformed habitats. This result sheds light on the way that biodiversity responds to anthropogenic transformations, showing higher stability than theoretically predicted.

## Introduction

Anthropogenic landscape transformations have caused the loss and fragmentation of many natural habitats with progressive and detrimental effects on species abundances and the continuous decrease of habitat quality and quantity (Fahrig et al. 2011). This is why forest loss due to anthropogenic landscape transformations is considered the single most pervasive threat to biodiversity worldwide (Fahrig 2013). However, more often than not, land use changes produce a mosaic of habitat patches with different quality and characteristics, where species may be able to use resources from patches of alternative and modified habitats in addition to resources usually found in patches of their original natural habitats (Brotons et al. 2005).

Species do not function as isolated entities, instead they are organized in complex networks of interspecific interactions (Palacio et al. 2016). A network is exemplified by plant– frugivore interactions in tropical forests, where fruit-eating vertebrates disperse the seeds of up to 90 percent of plants (Fleming et al. 1987). These networks substantiate the ecological processes that ultimately provide valuable ecosystem services for human societies (Bascompte and Jordano 2014). Therefore, if species’ occurrences and abundances decrease as a consequence of landscape modification, the interactions between species and the structure of the interactive networks is also expected to change (Woodward et al. 2010). By understanding what causes changes in ecological networks and the consequences these changes have on species interactions, it should be possible to predict how ecological processes and ecosystem services can respond to further landscape transformations (Valiente-Banuet et al. 2014).

The network structure may be related to the ecological systems’ emerging properties such as nestedness, modularity, and complementary specialization, which are key factors for the stability of ecological communities, the maintenance of ecological processes, and biodiversity conservation (Tylianakis et al. 2010). Nestedness measures how much (and how many) of the interactions among specialists species are a subset of the interactions among generalist species (Jordano et al. 2003). A nested structure minimizes competition and increases the number of coexisting species (Bastolla et al. 2009) and also makes the community more robust to both random extinctions (Burgos et al. 2007) and habitat loss (Fortuna and Bascompte 2006). Modularity characterizes the existence of densely connected, non-overlapping subsets of species (i.e. modules), which are composed of species having many interactions among themselves but with very few interactions between species from other modules (Castaño et al. 2018). Modularity increases network stability, retaining the impacts of a perturbation within a single module and minimizing impacts on other modules (Fortuna et al. 2010). Complementary specialization (H2′) measures the extent by which specialist species interact with other specialist species (Blüthgen et al. 2006). H2′ apprises us if there is high or low niche differentiation in the network. The importance of understanding why H2′ is changing has its basis in the fact that the system is losing species that are always specialists or because species behaving as specialists in that situation are being lost or becoming more generalist (Soares et al. 2017).

Not all species are equally important for the dynamics and stability of an ecological network (Martín González et al. 2010). Some species are central to the stability of the network; the structure of the network breaks down faster when central species are selectively removed compared to random removals of other species (e.g. Memmott et al. 2004). Centrality metrics are a useful tool for assessing the relative importance of species (Palacio et al. 2016). Different centrality indices measure different aspects related to the position of a species within its network. For example, closeness centrality (CC) measures the proximity of a node to all other nodes in the network (i.e. nodes with high CC values can rapidly affect other nodes and vice versa). Alternatively, betweeness centrality (BC) describes the importance of a node as a connector between different parts of the network. Nodes with BC > 0 connect areas of the network that would otherwise be sparsely or not connected at all (Martín González et al. 2010). Furthermore, species may be ranked according to each of these metrics as a way to choose target species for conservation efforts (Palacio et al. 2016). Such target species may also serve as proxies to evaluate changes due to habitat loss, fragmentation, and climate change (Nielsen and Totland 2014).

Even though it is expected that land use changes should modify network characteristics, there is still theoretical controversy and few empirical studies show how the loss of natural environments affect network nestedness, modularity, complementary specialization and centrality (Moreira et al. 2018). In this study, we investigated how a network of fruit-eating bats and their food sources responds to land use change in the Colombian Andes. We studied bat-fruit networks in three different habitats: continuous forests, forest fragments, and crops, across a gradient from pre-montane forest to a heterogeneous agricultural matrix known as the “Coffee Cultural Landscape of Colombia”. The “Coffee Cultural Landscape of Colombia” is a particular combination of coffee-growing cultural customs and sustainable practices that shows how farmers have adapted to the difficult geographic conditions of the area (Martínez Moreno 2011).

By analyzing bat–fruit networks from three habitats representing a gradient of modification we empirically validate whether the composition and structures of mutualistic networks are conserved after land use change in an active agricultural landscape. We ask the following questions: 1) Are network structural properties of bat–fruit networks conserved along a gradient of modification? 2) Do species’ roles within the bat–fruit network change along the transformation gradient? We hypothesize that the bat–fruit networks in all the studied habitats, despite contrasting modification history, will display nestedness and modularity, structural properties previously found in other mutualistic networks (Tylianakis et al. 2010).

## Methods

### Study area

The study area is located on the western slope of the Colombian Central Cordillera of the Andes in the municipalities of Santa Rosa de Cabal, Marsella and Dosquebradas (Department of Risaralda ca 4°54′N, 17°39′W). The study area ranges in elevation from 1616 m to 1990 m. UNESCO recognizes this region (including the departments of Risaralda, Quindío, Caldas and Valle del Cauca) as a World Heritage site known as the “Coffee Cultural Landscape of Colombia”. In this region coffee has been produced in small plots for more than 100 years and is one of the most important crops in the area (Martínez Moreno 2011). Other important crops are pastures, bananas, vegetables, forest plantations and other fruits (Villamil-Echeverri et al. 2015). The annual mean temperature oscillates between 16 - 24 °C; the mean relative humidity is 79% and the mean precipitation is 3358.4 mm (Cenicafé 2011, Jaramillo et al. 2011).

We selected nine sampling localities representing three habitats (3 localities for each habitat: (1) continuous forests, (2) forest fragments immersed in a matrix of crops, and (3) crops without forests). Each sampling locality was in the center of a 1 km buffer of each habitat; each sampling point was separated by at least 4 km from other sampling points. We follow a factorial design (Supplementary material Figure A1).

### Data sampling

We captured bats using mist nets (12 meters long) located at 1–5 m above the ground between August 2016 and August 2017. Each locality was surveyed four consecutive nights every three months. We used 5–7 mist nets (12×2.5m; 30mm mesh) per survey; we opened mist nets at 18:00 p.m. and closed them at 06:00 a.m. In the event of ongoing heavy rain, the nets were closed. We avoided surveys during full moon nights in order to prevent the influence of lunar phobia (Saldaña-Vázquez y Munguía-Rosas 2013). In total, we accumulated 47012 net hours of sampling effort (forests: 12297; fragments: 19116; crops: 18599). Species were identified using the most updated taxonomic key for the region (Diaz et al. 2016). Captured bats were held in cloth bags for no longer than 2h to allow them to defecate so we could maximize sample yield. We cleaned the bags thoroughly between captures to prevent cross-contamination of fecal samples. Bats were released after the collection of data and fecal samples. Voucher specimens were collected to represent the species diversity of bats at each sampling locality and were deposited in the “Colección de Vertebrados UNISARC (CUS-M 283-321)”. Each sample from every bat was collected separately and then dried and stored in plastic bags. Seeds were identified up to species level based on a reference collection of the study area (Rodríguez-Duque 2018). Seed samples were deposited in “Herbario UNISARC (CUS-P)”.

### Bat-fruit network structure

To analyze bat-fruit network structure we created binary and weighted matrices with bat species in rows and plant species in columns. Inside binary matrices, cell values of 1 (presence) represented an interaction between a specific bat species and a specific plant species. Zeros (absence) indicated no interaction. Weighted matrices were filled with the number of “consumption events”, defined as the sum of the presences of plant seeds in the fecal samples of each bat species (Castaño 2009). We created matrices for each of the nine study plots separately and for the three habitat types by pooling the data obtained from the three study plots within each habitat.

#### Species and interaction richness

For each of the nine study plot matrices we calculated plant network species richness (*P*), bat network species richness (*B*), network size (*S=B+P*) and interaction richness (*I=* cell values of 1 inside binary matrices.

#### Complex network metrics

For each of the three habitat type matrices (calculated by pooling the three study plots within each habitat) we calculated: (1) *Complementary Specialization* (H2′), varying from 0 (all bat species interacting with the plant species) to 1 (each species interacting with a particular subset of partners). This determined high or low niche differentiation in the network and could be used to compare which networks have more interactions between generalist or specialist species (Blüthgen et al., 2006). Then we calculated (2) *Weighted Nestedness* (WNODF) ranging from 0 (non-nested) to 100 (fully nested), to measure how strongly species interactions of seldom connected species were nested within those of highly connected species (Almeida-Neto and Ulrich 2011). To test whether estimates of WNODF differed significantly from networks with randomly interacting species, we compared the observed nestedness with the nestedness of 1000 random networks based upon a Patefield null model (Dormann et al. 2009). Finally we calculated (3) *Quantitative Modularity* (QuanBiMo) ranging from 0 (non-modular) to 100 (fully modular) using the algorithm QuanBiMo (Dormann and Strauss 2014). This algorithm uses the hierarchical random graph approach, which organizes interacting species into a graph so that close species are more likely to interact. Then it swaps branches randomly at different levels and reassesses the modularity of the network selecting the more modular organization QuanBiMo. To test the significance of modularity, we generated 1000 random networks fixing the probability that two species would interact based on the observed real networks.

We used the Patefield null model to estimate the significance of the observed network metrics. We then calculated the modularity of the networks and evaluated whether observed modularity fell within the 95% confidence interval calculated from the randomized matrices. We finally standardized the modularity by calculating the Z-score Q (ZQ).

#### Species roles within the networks

In each of the habitat type matrices we assessed the functional role of bat and plant species using three centrality metrics. The (1) *degree centrality* (DC) measured the number of interactions of a given species, reflecting its degree of generalization versus specialization, The (2) *betweenness centrality* (BC) measured the extent by which a species acts as a connector to the lowest number of direct or indirect interactions among other pairs of species, The (3) *closeness centrality* (CC) was the mean of the lowest number of direct or indirect interactions from one species to every other species in the network with higher numbers yielding lower CC scores (Palacio et al. 2016).

### Comparison between landscapes

To compare if there were any effects of habitat modification on the species and interaction richness in the network we made post hoc comparisons using a Duncan test. Variables were log transformed to meet assumptions of the tests. In order to compare if there were any effects of landscape modification on the complex network metrics (H2′, WNODF and Q) we compared pairs of networks using monte carlo procedures (Muylaert and Dodonov 2016). To test whether a particular species’ functional role within the networks varied among the three habitat types we used the Pearson correlation coefficient for each centrality metric (DC, BC, CC) between pairs of habitats. Our goal was to assess whether the functional role of a species in one habitat could explain the same species functional role in another habitat. In these analyses, we pooled the three networks over three study plots rather than the nine separate networks as the number of species found in all nine networks was smaller.

## Results

We captured thirteen bats species (Phyllostomidae, subfamilies Stenodermatinae and Carolliinae) that consumed 37 plant species belonging to 11 botanical families (Araceae, Campanulaceae, Cyclanthaceae, Ericaceae, Gesneriaceae, Hypericaceae, Moraceae, Myrtaceae, Piperaceae, Solanaceae y Urticaceae) (Supplementary material Table A1). The plants most consumed by bats were *Cecropia telealba* (Urticaceae, 22% of all fecal samples), *Solanum aphiodendrum* (Solanaceae, 17%), *Ficus americana* (Moraceae, 9%), *Piper aduncum* (Piperaceae, 6%), *Piper crassinervium* (Piperaceae, 6%), and *Ficus tonduzi* (Moraceae, 5%).

The continuous forest habitat contained the lowest network size, interaction richness, and richness of plants on average. The bat network species richness was not different between habitats (Figures 1 and 2). The three networks (each habitat) were all nested, modular and specialized compared with respective null models (Supplementary material Table A2). Moreover, there were no differences between landscapes in nestedness or modularity; however, the crop habitat contained less specialized networks compared to continuous forests and fragments (Figure 3).

**Figure 1.**
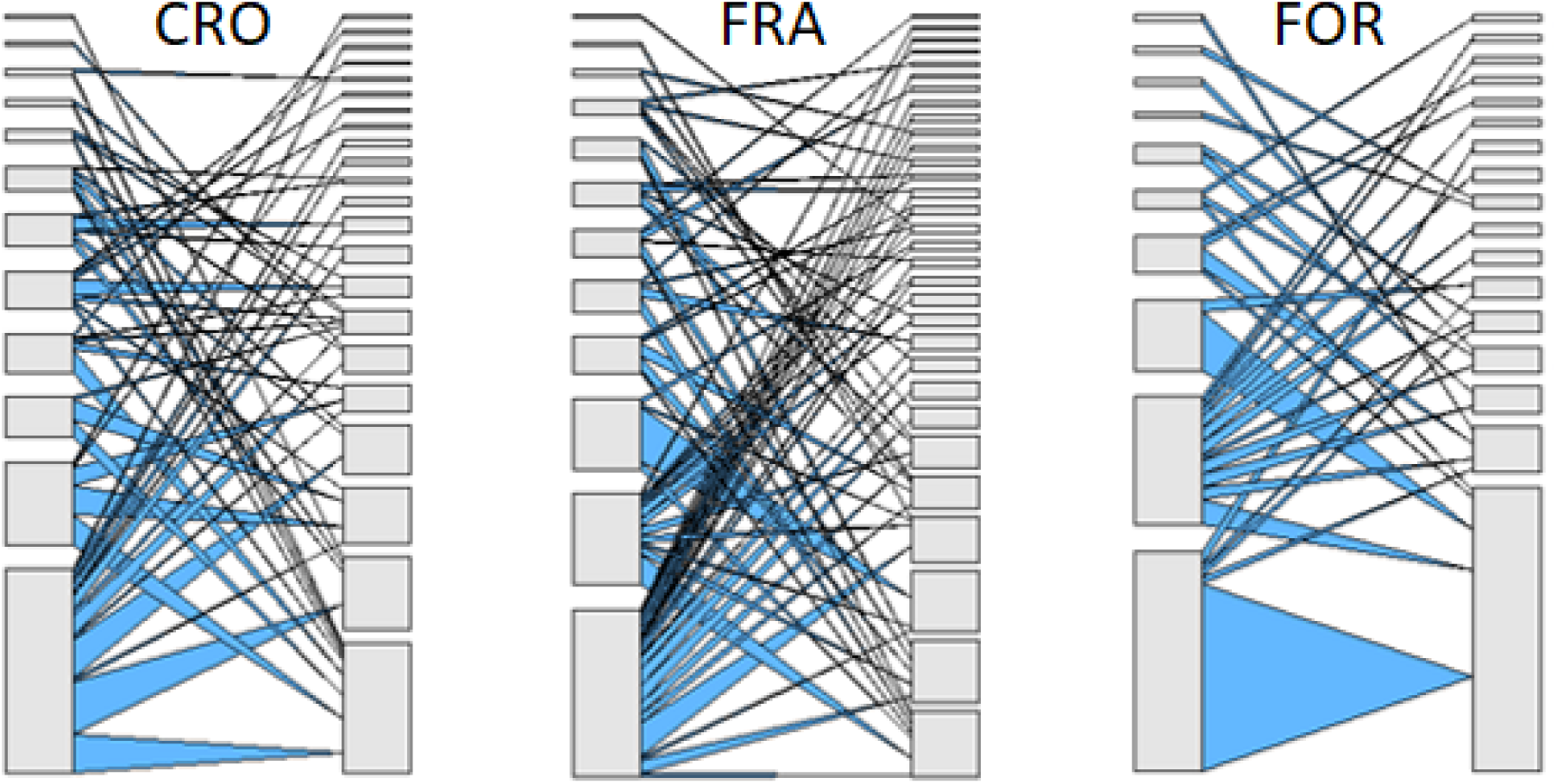
Quantitative bat–fruit networks along a gradient of increasing habitat modification. For each web, left bars represent bat abundance and right bars represent fruit plant abundance, drawn at different scales. Linkage width indicates frequency of each trophic interaction.

**Figure 2.**
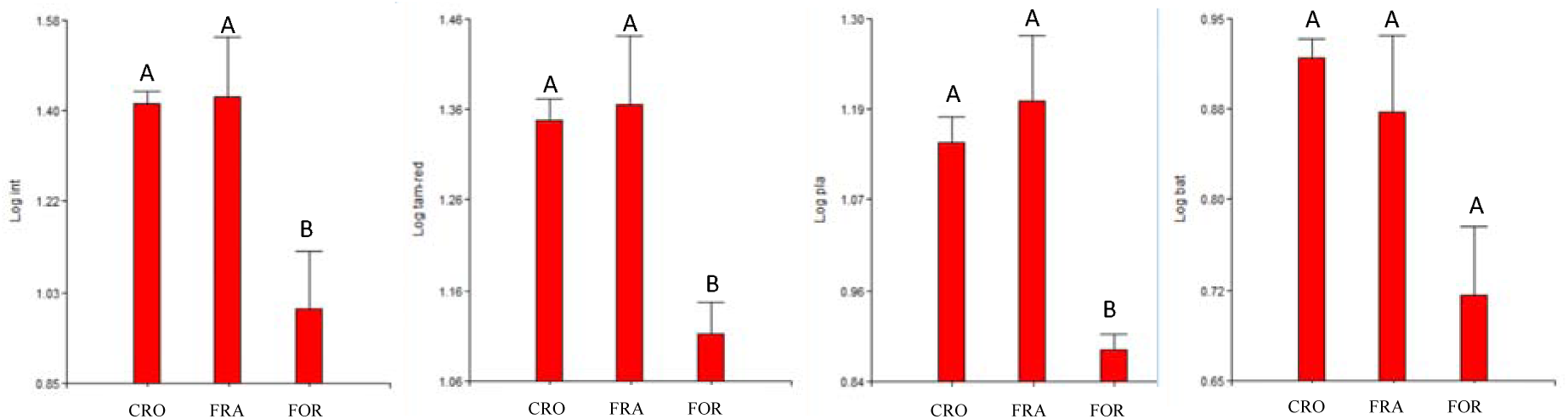
The effects of habitat modification on four qualitative network metrics (plant richness, bat richness, network size and interaction richness). Whiskers represents standard error, letters above individual means indicate significant differences (P≥0.05) among habitat types (FOR:continuous forests, FRA: forest fragments and CRO: crops) for that particular metric, letters in common indicates no significant difference.

**Figure 3.**
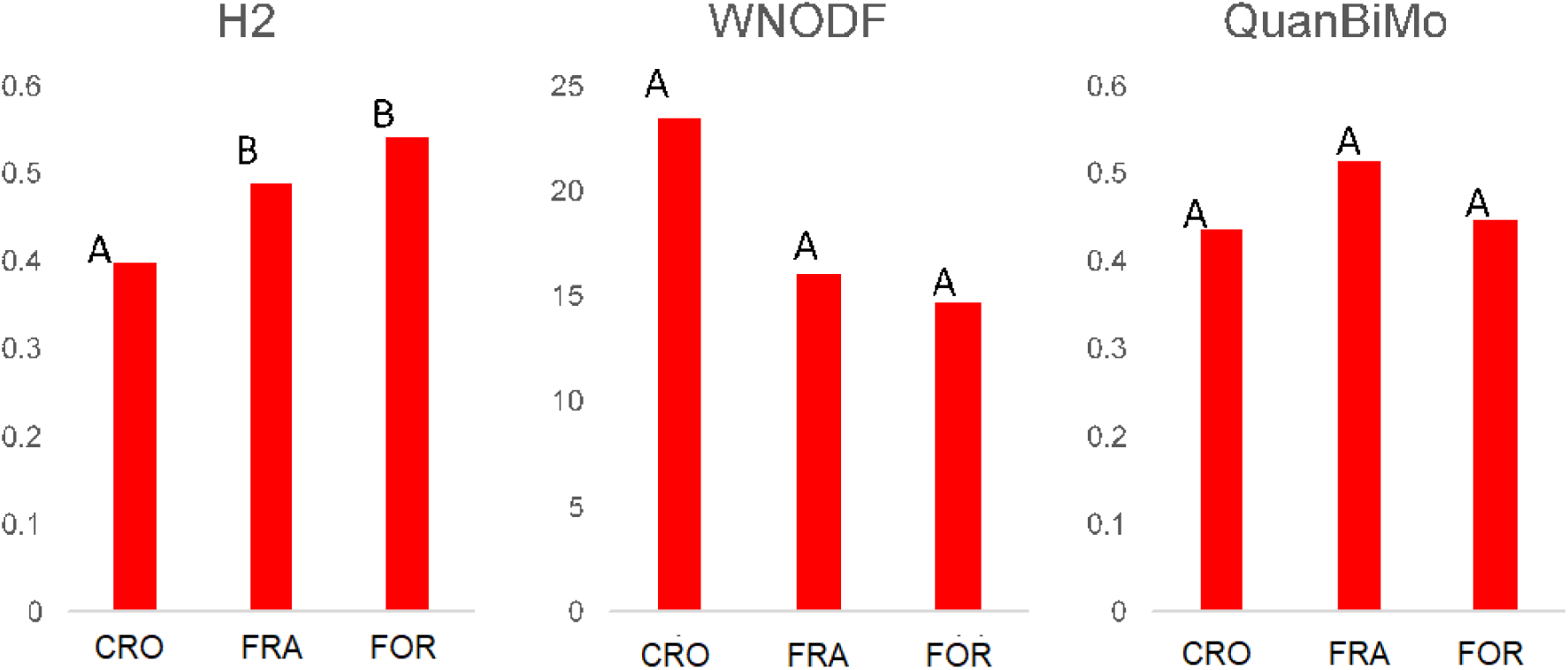
The effects of habitat modification on three quantitative complex network metrics (H2′, WNODF and QuanBiMo). **H2**′ Complementary Specialization varies from 0 (all species interacting with the same partners) to 1 (each species interacts with a particular subset of partners). **WNODF** Weighted Nestedness ranges from 0 (non-nested) to 100 (fully nested). **QuanBiMo** Quantitative Modularity ranges from 0 (non-modular) to 100 (fully modular). Letters indicate significant differences (P≥0.05) among habitat types (FOR:continuous forests, FRA: forest fragments and CRO: crops) for that particular metric, letters in common indicates no significant difference. We compare pairs of networks using Monte Carlo procedures (Muylaert and Dodonov 2016).

### Species’ roles within the network across habitats

The most central bat in continuous forest and fragments was *Carollia brevicauda*, whereas in crops it was *Artibeus lituratus*. Similarly the most central plant species in forest and fragments was *C. telealba*, whereas in crops it was *S. aphiodendrum* (Supplementary material Table A3).

There was no pairwise correlation in DC, BC and CC for particular plant species between crops and the other two habitats (forest fragment and continuous forest). For bats, however, the DC (number of interacting partners) showed low correlation (R^**2=**^ 0.39, *P < 0.05*) between crop and forest fragments. There was no pairwise correlation in BC and CC between crops and the other two habitats. A negative correlation would have indicated a systematic change, but the lack of correlation suggests a more random pattern. The pair-wise correlations between forest fragments and continuous forest were significant (R^**2**^ = 48%-94%, *P < 0.*05) in all centrality metrics for both plant and bat species (Table1).

## Discussion

By analyzing the bat–fruit networks along a gradient of increasing habitat modification in the “Coffee Cultural Landscape of Colombia” we observed the following. The non-modified habitat (continuous forest) contained smaller bat–fruit networks than the modified forest fragments and crops. The modified landscapes had similar ecological network structures compared to continuous forest, with no significant differences observed for nestedness and modularity metrics. Despite similar network properties among the three levels of habitat modification, the more modified habitats induced a homogenization of the bat–fruit networks; crops contained less specialized networks and the species’ roles in crops changed in relation to the roles in continuous forests and fragments.

### Species and richness of interaction

The non-modified habitat (continuous forest) contained, on average, smaller network size, lower interaction richness, and lower plant richness than did forest fragments and crops. Although no significant differences in bat richness were detected among habitats, continuous forest had a slightly lower bat richness than the modified habitats. Although the effect of landscape transformations on seed dispersal networks has not been evaluated hitherto (Hagen et al. 2012), this result is consistent with previous studies on plant-pollinator networks where forests contained smaller plant–pollinator networks than the more disturbed habitats (Hagen and Kraemer 2010, Nielsen and Totland 2014). These results may indicate that some agricultural landscapes are not hostile for bat and plant species; in fact, it can harbor complementary resources (*sensu* Brotons et al., 2005) favoring interactions that are typical of both modified and unmodified habitats.

### Complex network metrics

Modified habitats (e.g. forest fragments and crops) had similar bat-fruit network structures to continuous forest with no significant differences observed for nestedness and modularity metrics. Few empirical studies have shown that structural properties of ecological networks withstand habitat degradation (Tylianakis et al. 2007, Nielsen and Totland 2014). However, theoretical models have suggested that networks can remain robust as long as habitat loss does not exceed 80%. Below this threshold, networks showed massive and rapid species extinctions (Fortuna and Bascompte 2006). The fact that the bat-plant networks maintain their structure in this agricultural landscape and the fact that species and interactive richness were high even in habitats with a high degree of modification suggests that forest loss in this type of agricultural system does not necessarily lead to a cascade of secondary extinctions that would lead to an ecosystem collapse. Rather, new interactions are reconfigured where bats feed on plants that grow in the crops (e.g. *Psidium guajava* or *Solanum aphyodendrum*). This result sheds light on the way that biodiversity can respond to particular anthropogenic transformations, showing higher stability than theoretically predicted. This is consistent with previous studies that have shown that the heterogeneity of the “Coffee Cultural Landscape of Colombia” is able to maintain high levels of biodiversity that provide valuable ecosystem services (Carranza-Quiceno et al. 2018). Thus, appropriately managed agroecosytems can retain ecological networks that are structurally and functionally similar to unmodified habitats (Tylianakis et al. 2007). Nonetheless, we must take these results with caution because, despite similar network properties among the three levels of habitat modification, the more modified a habitat is, the more homogeneous its bat–fruit networks become. For example, the crop habitat contained less specialized networks and the species role in this habitat changed, compared to those in continuous forests and fragments.

### Species role within the network in three landscapes

The role of bat and plant species in continuous forest was correlated with their role in forest fragments. In contrast, the species roles in crops did not correlate with the other habitats. This means that both plants and bats fulfill similar roles in the less transformed habitats that retain some forest cover, but they change their roles in crops that represent the highest transformation of the study area landscape, in this case open areas with isolated trees and shrub vegetation predominate.

With regards to bats, *C. brevicauda* has been considered the most important montane frugivore bat both in Central and South América in terms of richness of plants consumed (Castaño et al. 2018). In this study *C. brevicauda* was the central-most bat in forests and fragments, however in crops it was replaced by *Artibeus lituratus*. This switch in centrality suggests that functional traits could be influencing the species′ role in plant-frugivore networks. For example, *A. lituratus* is a larger and heavier bat and probably faces lower predation risk compared to other frugivores (Cohen et al. 1993), and these traits seem more suitable for occupying modified landscapes (Saldaña-Vázquez and Schondube 2015). The smaller but more maneuverable *C. brevicauda* would favor flight in spatially complex environments such as the interior of forests and fragments of forest (Marinaello and Bernard 2014).

Regarding plants, *C. telealba* is a tall tree, and was considered the most consumed plant by bats in a subandean landscapes (Aguilar-Garavito et al. 2014). In our study we also found that *C. telealba* was the most central plant in forest and fragments, however in crops it was replaced by a bush *S. aphyodendrum*, the latter, having a smaller size and faster maduration. which would be more likely to inhabit highly changing enviroments rather than crops (Falster et al. 2018).

Besides our suggestion that functional traits could be influencing the species role in plant-frugivore networks, several studies have found that different ecological and physical attributes such as animal body mass (Chamberlain and Holland 2009, Sebastián-gonzález et al. 2017), dietary specialization (Mello et al. 2015), seed width, and length of plant fruiting period (Vidal et al. 2014) explain the structural importance of the species in mutualistic networks. However, this is the first study with evidence of changes in the species role within interaction networks in response to habitat transformation (Nielsen and Totland 2014). Future studies should evaluate which variables affect changes in the role played by species in interaction networks.

## Conclusion

As far as we are aware, this is the first study that evaluates the response of bat-fruit networks to a gradient of habitat transformation. Our results suggest that the networks in the “Coffee Cultural Landscape of Colombia” maintain their structure across habitats in a landscape with different land uses, which indicates that the seed dispersal service is maintained even in the most transformed habitat. High network richness could be related to the high heterogeneity present in this agroecosystem. Future studies should evaluate how landscape heterogeneity affects interaction networks. Our results have important implications for the conservation of biodiversity and the maintenance of ecosystem services-particularly seed dispersal - because we have documented the most central and important plant and animal species in the network. Even though the number of species does not decrease due to habitat transformation, species change their role in the most transformed habitat.

**Table 1.** Pair-wise correlation tests (Pearson’s correlation) between the three landscape types (continuous forests, forest fragments and crops) based on three centrality metrics (DC,BC,CC). **DC** Degree Centrality indicates the number of interacting species of all the species occurring in all three landscapes sampled in Coffee Landscapes of Colombia. **BC** Betweeness Centrality measures the extent to which a species acts as a connector on the lowest number of direct or indirect interactions among other pairs of species. **CC** Closeness Centrality is the mean of the lowest number of direct or indirect interactions from one species to every other species in the network. (*) = *P* < 0.05, (**) = *P* <0.01.

| Landscapes | Dc |  | Bc |  | Cc |  |
| --- | --- | --- | --- | --- | --- | --- |
|  | R<sup>2</sup> Bat | R<sup>2</sup> Pla | R<sup>2</sup> Bat | R<sup>2</sup> Pla | R<sup>2</sup> Bat | R<sup>2</sup> Pla |
| For-Fra | 0.94** | 0.48** | 0.94** | 0.69** | 0.57* | 0.16 |
| Fra-Cro | 0.39* | 0.1 | 0.36 | 0.09 | 0.12 | 0.07 |
| Cro-For | 0.17 | 0.18 | 0.16 | 0.23 | 0.22 | 0.05 |

## Declarations

We thank M. Vélez, P Vélez, L.M. Romero, D.L. Rodríguez, C. Villabona, and C. López for their assistance in the field. This work was supported by the Patrimonio Autónomo Fondo Nacional de Financiamiento para la Ciencia, la Tecnología y la Innovación Francisco José de Caldas [code 643771451270]; and the Vicerrectoría académica Pontificia Universidad Javeriana [code 1215007010110]. All authors formulated the idea: the first and second author conducted the fieldwork, first author performed analyses, all authors wrote the manuscript. No potential conflict of interest was reported by the authors. Corporación Autónoma Regional de Risaralda approved animal’s capture (CARDER-, license number 2004-Sep 2016) and the Animal Use and Care Committee of the Corporación Universitaria Santa Rosa de Cabal-UNISARC approved all the procedures.

