## Supplementary material for "Bat-fruit networks structure resist habitat modification but species roles change in the most transformed habitats"

**Supplementary material Figure A1**. Study area with nine sampling localities representing three habitats: continuous forests (FOR), forest fragments immersed in a matrix of crops (FRA), and crops without forests (CRO). We follow a factorial design with three levels of habitat modification and three levels of elevation.

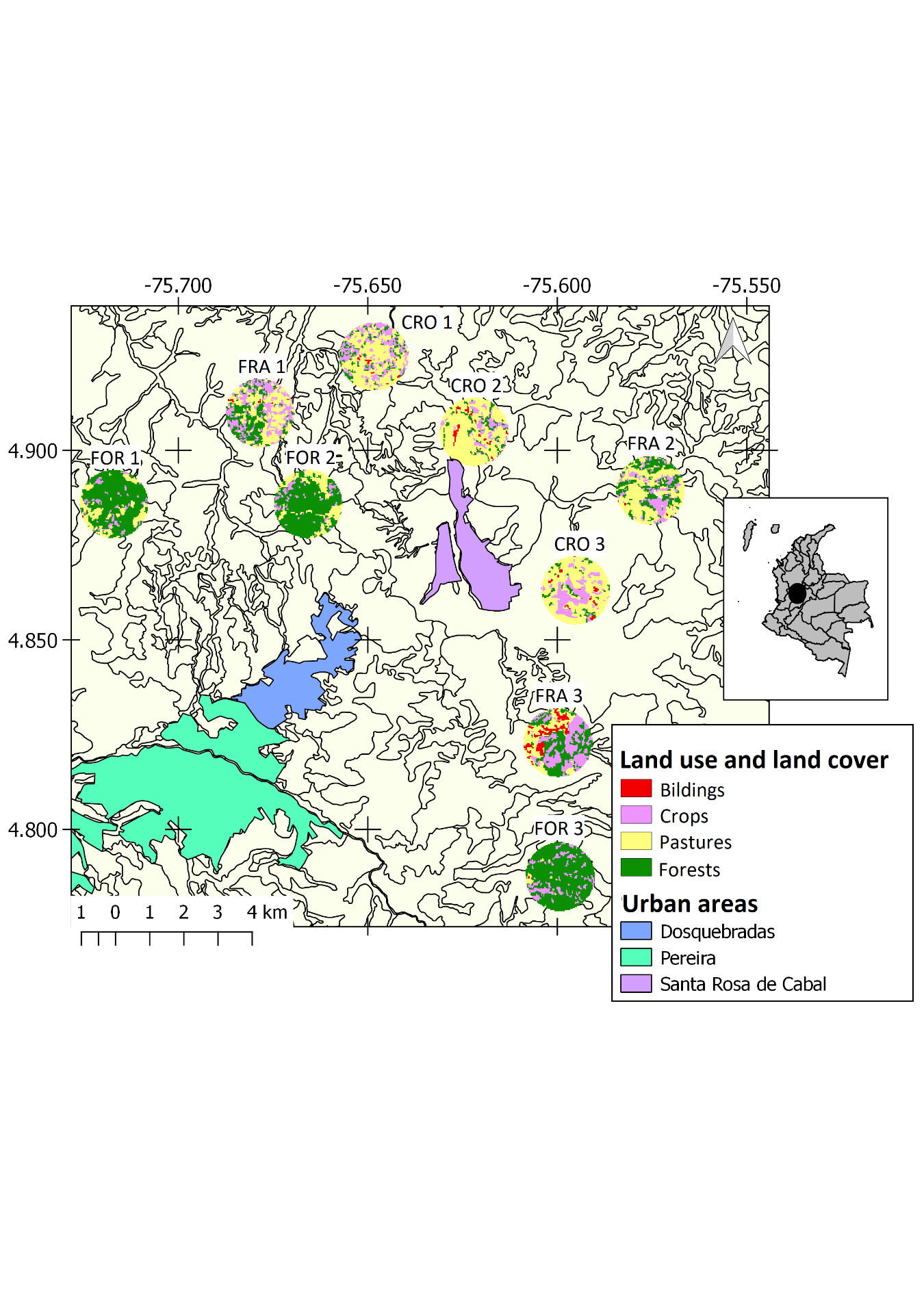

|  |  | **Gradient of increasing habitat modification** | | |
| --- | --- | --- | --- | --- |
|  |  | **FOR** | **FRA** | **CRO** |
| **Gradient of increasing elevation** | **1** | FOR 1: 1646m. | FRA 1: 1690m. | CRO 1: 1616m |
|  | **2** | FOR 2: 1831m. | FRA 2: 1820m | CRO 2: 1792m. |
|  | **3** | FOR 3: 1930m. | FRA 3: 1990m. | CRO 3: 1919m. |

**Supplementary material Table A1.** General interaction matrix describing plant species consumed by Phyllostomid bats in the study area.

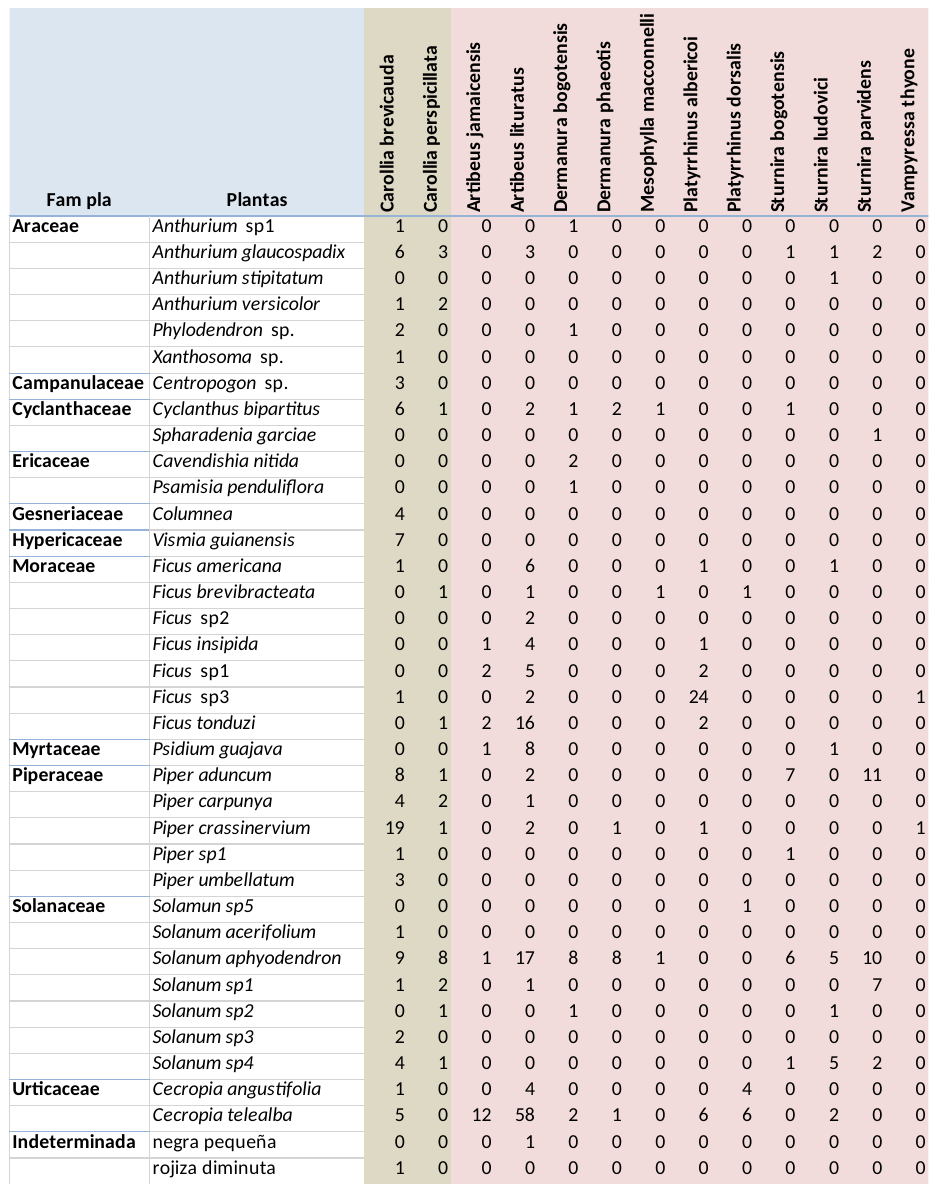

**Supplementary material Table A2.** Qualitative measures of the three bat–fruit networks studied in Colombian Coffee Cultural Landscape. ‘CRO, FRA and BOS represents the pooled bat–fruit networks from three sample plots. H2 is the Complementary Specialization index. WNODF is the nestedness index. We compared the observed nestedness with the nestedness of 1000 random networks based upon a Patefield null model. QuanbiMo is the modularity index calculated. We compared the observed modularity with the modularity of 1000 random networks based upon a Patefield null model Three asterix (***) behind the value indicates that the value is significantly higher than would be expected by chance.
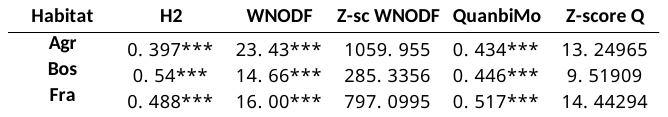

**Supplementary material table A3** Centrality metrics (Dc, Bc, Cc) per bat and plant species in the three landscape types sampled in Colombian Coffee Cultural Landscape . Degree centrality (DC) Betweeness centrality (BC) and Closeness centrality (CC). Abbreviation of the bat and plant species correspond to firs letter of the genus and three first letters of the species, full names in Supplementary material Table A1
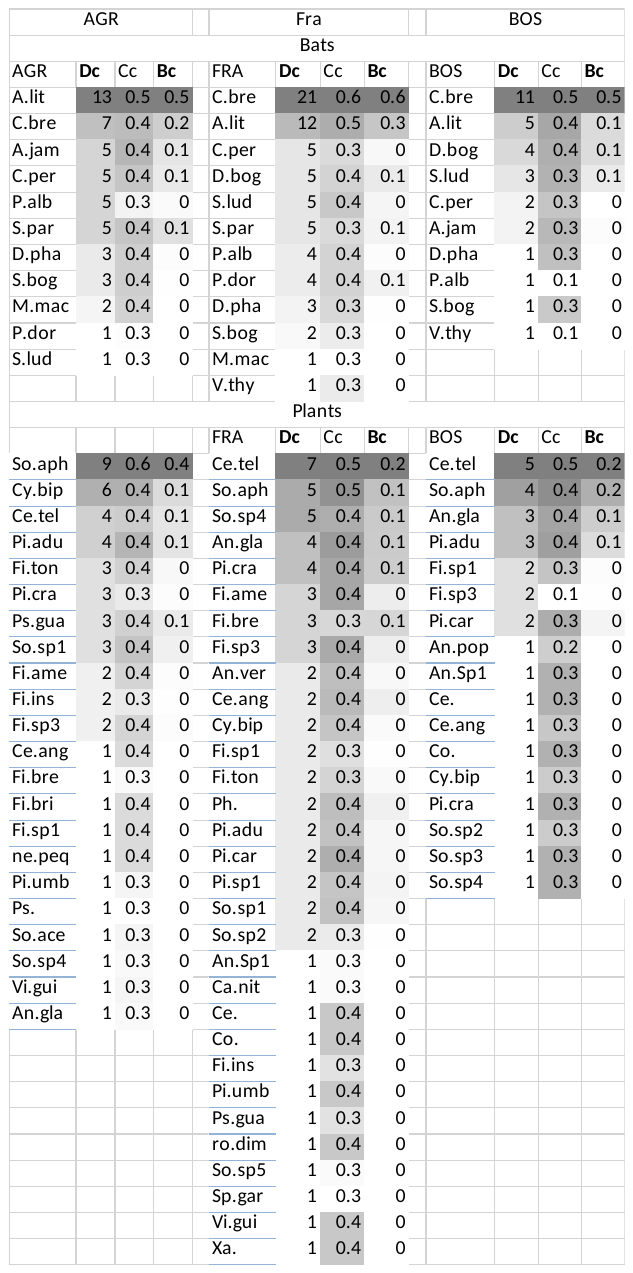
